# Origin of gibberellin-dependent transcriptional regulation by molecular exploitation of a transactivation domain in DELLA proteins

**DOI:** 10.1101/398883

**Authors:** Jorge Hernández-García, Asier Briones-Moreno, Renaud Dumas, Miguel A. Blázquez

## Abstract

DELLA proteins are land-plant specific transcriptional regulators known to interact through their C-terminal GRAS domain with over 150 transcription factors in *Arabidopsis thaliana*. Besides, DELLAs from vascular plants can interact through the N-terminal domain with the gibberellin receptor encoded by *GID1*, through which gibberellins promote DELLA degradation. However, this regulation is absent in non-vascular land plants, which lack active gibberellins or a proper GID1 receptor. Current knowledge indicates that DELLAs are important pieces of the signalling machinery of vascular plants, especially angiosperms, but nothing is known about DELLA function during early land plant evolution or if they exist at all in charophytan algae. We have now elucidated the evolutionary origin of DELLA proteins, showing that algal GRAS proteins are monophyletic and evolved independently from those of land plants, which explains why there are no DELLAs outside land plants. *DELLA* genes have been maintained throughout land plant evolution with only two major duplication events kept among plants. Furthermore, we show that the features needed for DELLA interaction with the receptor were already present in the ancestor of all land plants, and propose that these DELLA N-terminal motifs have been tightly conserved in non-vascular land plants for their function in transcriptional co-activation, which allowed subsequent exaptation for the interaction with the GID1 receptor when vascular plants developed gibberellin synthesis and the corresponding perception module.

## Introduction

DELLA proteins are transcriptional regulators that have been extensively characterized during the past twenty years (Vera-Sirera et al. 2015). They are involved in diverse processes ranging from seed germination to flowering, including legume nodulation, stress responses, or fern sexual reproduction (Peng and Harberd 1997; Floss et al. 2013; Tanaka et al. 2014). In fact, these proteins are responsible for the dwarf phenotype that allowed the development of new crop varieties during the Green Revolution (Peng et al. 1999). DELLAs are one of the main elements that compose the gibberellin (GA) signalling pathway in vascular plants, acting as the negative regulators of the pathway (Dill et al. 2001; Itoh et al. 2002).

As part of the GRAS family of plant specific proteins, they present a highly conserved C-terminal domain, the GRAS domain. Initially, this domain was suggested to be distantly related to the STAT family of metazoan proteins (Richards et al. 2000). More recently, a thorough *in silico* structural analysis of the domain has evidenced a remarkable similarity to bacterial Rossman-fold SAM-dependent methyltransferases suggesting a bacterial origin of the GRAS domain (Zhang et al. 2012). Even though no chlorophytan alga presents GRAS-like genes, several charophytan species contain genes encoding GRAS proteins, pointing to an streptophytan origin of the family (Engstrom 2011; Delaux et al. 2015).

The GRAS domain of DELLA proteins is responsible for the establishment of protein-protein interactions. DELLAs cannot bind DNA, but they can interact with over 150 transcription factors and other transcriptional regulators, and modulate their functions in order to regulate gene expression (Marín-de la Rosa et al. 2014). DELLAs can either negatively affect transcription factor function, mainly through a sequestering mechanism, or positively enhance their ability to activate transcriptional activity (Locascio et al. 2013). This allows DELLAs to coordinate multiple transcriptional programs and may have been an important trait acquired during plant evolution (Briones-Moreno et al. 2017).

A second important characteristic of DELLA proteins is their GA-dependent stability. This ability relies in the N-terminal, DELLA domain. The gibberellin receptor GIBBERELLIN INSENSITIVE1 (GID1) is able to interact directly with this N-terminal domain after GA binding (Ueguchi-Tanaka et al. 2005; Griffiths et al. 2006). Upon GA-GID1-DELLA complex formation, the SLY1/GID2 F-box protein interacts through the GRAS domain and recruits an SCF E3 ubiquitin ligase complex that marks DELLAs for degradation (Mcginnis et al. 2003; Sasaki et al. 2003; Dill 2004; Gomi et al. 2004). Three important motifs are involved in the interaction with GID1 proteins: DELLA, LEQLE and VHYNP. Mutations in these motifs impair GID1-DELLA interaction, giving rise to GA-resistant DELLA versions (Dill et al. 2001; Murase et al. 2008).

Every land plant genome sequenced so far contains *DELLA* genes, but their characteristic features related to GA signalling (i.e. N-terminal motifs and GA regulation, **Fig. 1A**) have been reported only in vascular plants (Hirano et al. 2007; Yasumura et al. 2007). These early studies indicate that GA-dependent DELLA regulation first appeared in the vascular plants common ancestor, however the analyses were constrained by the limited availability of genomic and transcriptomic resources from only the moss *Physcomitrella patens*, and the lycophytes *Selaginella moellendorffii* and *S. lepidophylla*. In these studies, neither a clear set of late GA synthesis genes nor proper GID1 receptors were detected in non-vascular plants, supporting the idea of GA-mediated regulation of DELLA proteins being vascular plant specific. In fact, no reports are available for the presence of bioactive GAs in mosses, and application of these compounds have no effect on moss growth (Hayashi et al. 2010).

**Figure 1.**
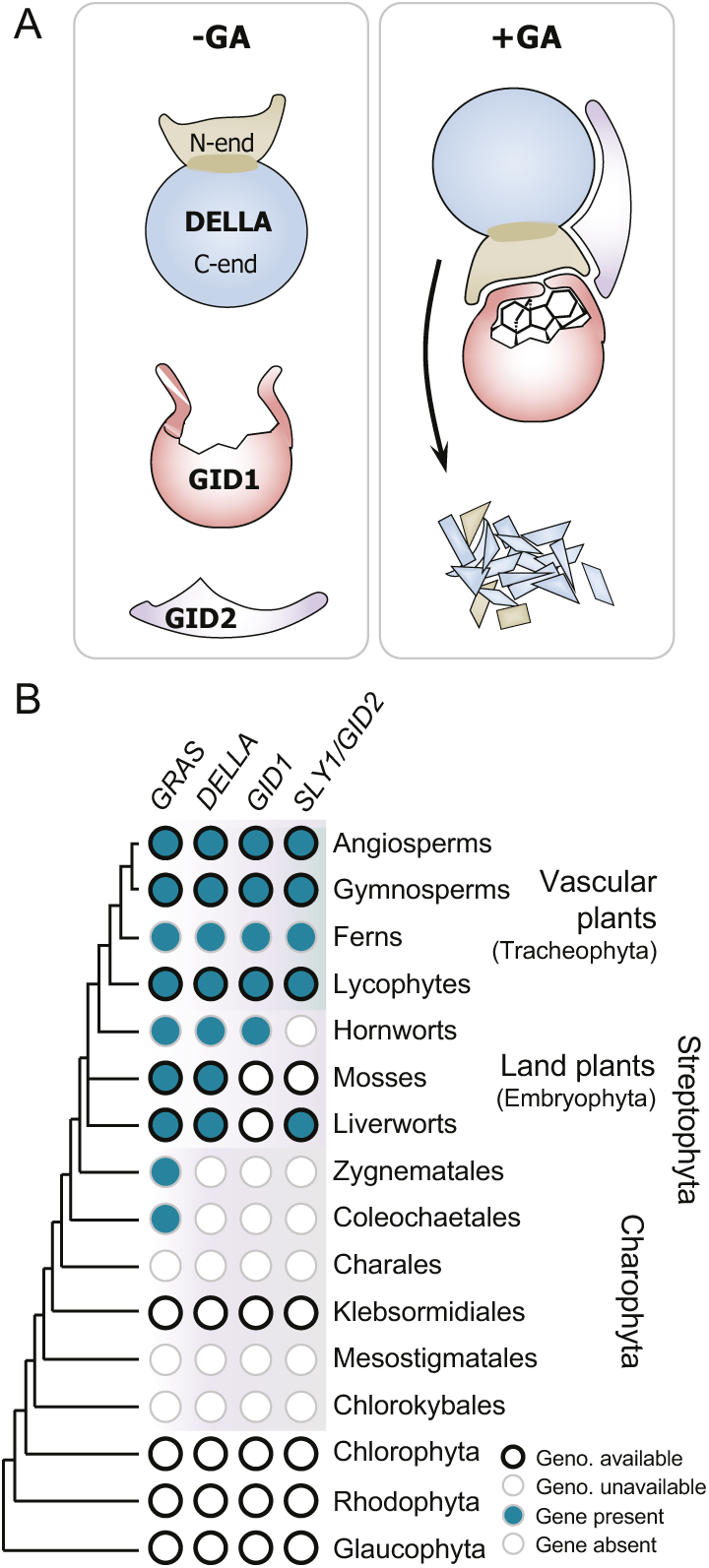
Gibberellin signaling elements are present in vascular plants. (*A*) Gibberellin signaling in vascular plants. (*B*) Presence of gibberellin-signaling related sequences in different phyla. GRAS, GID1 and GID2 orthologs were retrieved from oneKP or genome databases by BLASTP or TBLASTX searches. GRAS proteins were validated by positive Pfam (PF03514.13) and DELLA were counted either when Pfam (PF12041.7) was positive or BLASTP e-value was lower than 1E-20 when using either AtRGA or PpDELLAa. GID1 were counted by the presence described N-lid residue presence and α/β-hydrolase active site conservation. SLY1/GID2 were selected by thresholding BLAST results with 1E-20 based on similarity to *S. moellendorffii* GID2 proteins.

Current knowledge indicates that DELLAs are important pieces of the signalling machinery of late diverging land plants, especially angiosperms, but nothing is known about DELLA function during early land plant evolution or if they exist at all in charophytan algae. The previous lack of data can now be completed with new genomic and transcriptomic sources from earlier diverging land plants and algae to understand the origin and emergence of DELLA proteins. In fact, the recently sequenced genome of the liverwort *Marchantia polymorpha* encodes a DELLA protein whose N-terminal motifs are more similar to its vascular orthologs than to moss sequences (Bowman et al. 2017). In the present work we have tried to elucidate the evolutionary origin of DELLA proteins, and also investigate the functionality of the N-terminal domain by analysing the conservation and diversification of specific motifs in that domain. We found that algal GRAS proteins are monophyletic and evolved independently from those of land plants, indicating that there are no DELLAs outside land plants, and propose that the ancestral role of the N-terminal domain was as a transcriptional activation module which conservation allowed the co-option for the interaction with the GID1 receptor later during land plant evolution.

## Results

### Identification of GRAS and GA signalling elements sequences in plants

Previous studies have shown that GRAS domain sequences are present in several zygnematalean algae (Engstrom 2011; Delaux et al. 2015). We used these GRAS domains as bait to analyse available transcriptomes and genomes of all plants (i.e. Archaeplastida) in order to retrieve *GRAS* genes (**Fig. 1B**, **Supplementary Table 1**). After curation, we obtained GRAS sequences belonging to land plants and two groups of charophytan algae: Zygnematales and Coleochaetales. We did not find GRAS sequences in other algal groups, including the rest of charophytan groups, chlorophytes, rhodophytes and glaucophytes. Among GRAS sequences, we detected *bona fide* DELLA sequence hits in all land plant extant clades. We also searched for other known GA signalling components: the receptor GID1 and the F-box protein SLY1/GID2. Although we found similar sequences to GID1 in many clades (i.e. GID1-like proteins, or GLPs, Supplementary **Table 2**), those present in non-vascular plants do not contain the amino-lid sequences necessary for GA perception and DELLA interaction, and resemble those found in *Physcomitrella patens* (Hirano et al. 2007; Yasumura et al. 2007). Hence, we did not consider them as GID1 receptors. However, we found two hornwort sequences (an almost complete sequence for *Phaeoceros carolinianus* and a partial sequence for *Paraphymatoceros halli*) that represent good candidates for an ancestral state of GID1 receptors, pointing to a possible pre-tracheophytan origin of putative GID1 proteins (**Supplementary Fig. 1**). Among non-vascular land plants, we confirmed that the presence of SLY1/GID2 orthologous sequences is not only evident in *M. polymorpha* (Bowman et al. 2017), but in all liverworts examined, and absent in other non-vascular plants (**Supplementary Fig. 2**, **Supplementary Table 3**). These data consolidate the presence of GRAS in charophytan algae, not only in Zygnematales but also in Coleochaetales, and is consistent with the idea of GRAS proteins appearing first in charophytes before land colonization. Besides, DELLA proteins most likely appeared in the land plant common ancestor, but the GA signalosome components can only be found simultaneously in tracheophytes (**Fig. 1**). However, these views may change with further improvement of genomic data quality and availability from Charales and/or hornworts.

### Phylogenetic analysis of GRAS proteins

To elucidate the origin of the DELLA subfamily, we analysed the phylogenetic relationship between GRAS sequences in algae and land plants. For this, we used the GRAS domain of the sequences found, and added previously described eubacterial GRAS sequences to use as outgroup in a phylogenetic tree (Zhang et al. 2012). Interestingly, algal and land plant sequences formed two independent and statistically supported clades (**Fig. 2A**). This suggests that all land plant *GRAS* genes arose from a single gene present in an algal and land plant common ancestor. Further expansion and loss of GRAS subfamilies has occurred independently several times during plant evolution, and no clear correlation between the number of GRAS sequences and factors such as biological complexity seems to exist (**Fig. 2B**). Among land plants, we found sequences from twelve known GRAS subfamilies in *Arabidopsis thaliana* (SCL9, SHR, PAT1, SCL16/32, SCL29, SCL3, DELLA, SCL28, SCR, LAS, SCL4/7 and HAM), and sequences with no clear *A. thaliana* match in at least two early diverging land plants that resemble RAM1 sequences (**Fig. 2C**). We conducted phylogenetic analysis of these GRAS domain sequences and obtained highly supported clades for these groups in all land plant lineages (**Fig. 2D**). In fact, these groups greatly coincide with those recently published in the *Marchantia polymorpha* genome (Bowman et al. 2017). Altogether, these analyses indicate that previously known GRAS subfamilies are land plant specific and appeared early in a land plant common ancestor. Consequently, we consider that only land plant genomes may contain *DELLA* genes.

**Figure 2.**
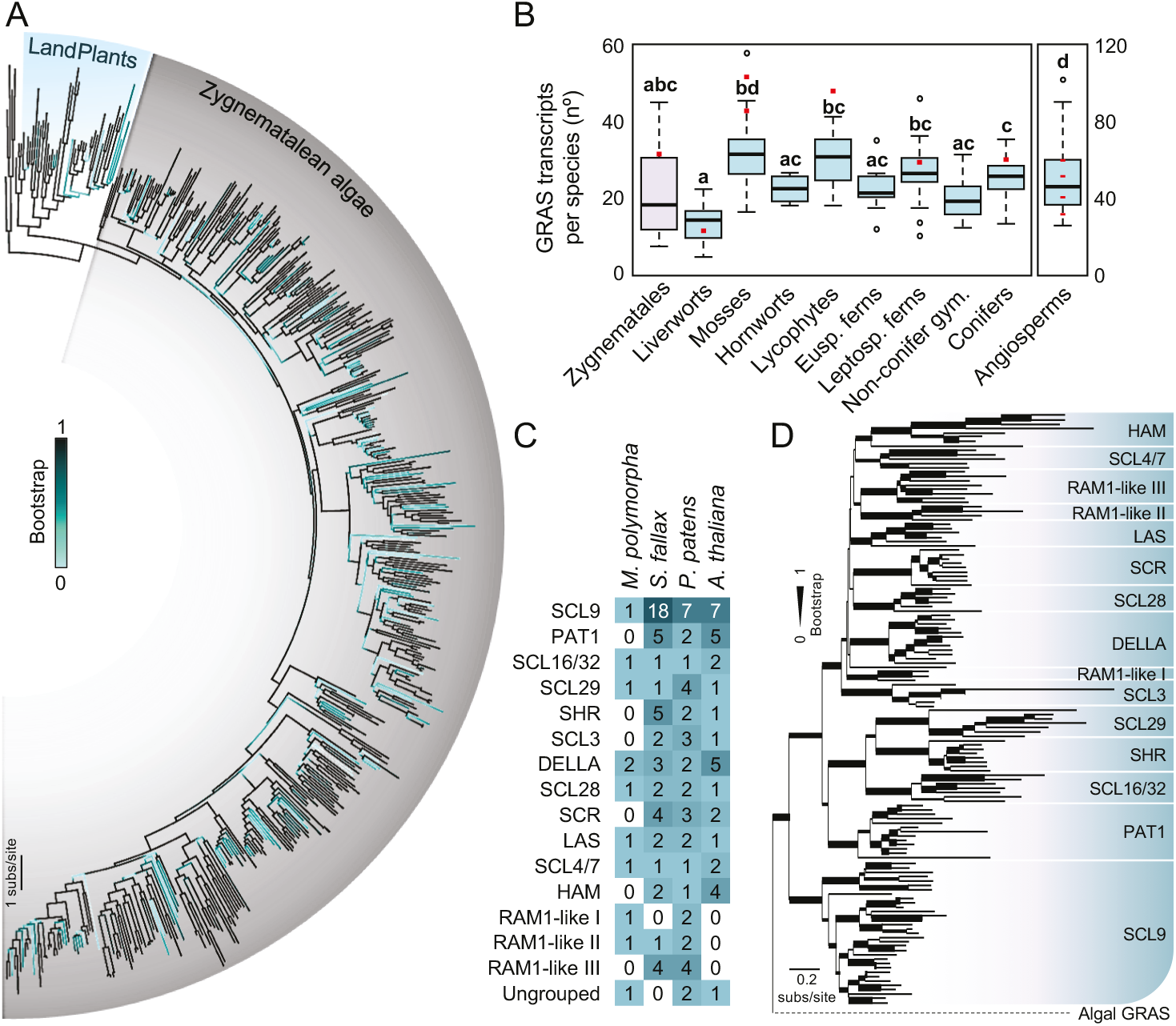
DELLA proteins are land plant specific. (*A*) Phylogenetic analysis of streptophytan GRAS proteins using GRAS domains. Support values associated with branches are maximum likelihood bootstrap values from 1000 replicates depicted as a color range (light turquoise to black). Black branches indicate a bootstrap of 1 (100%). Blue background denotes land plant (i.e. embryophytan) clade, grey background denotes algae sequences. (*B*) Number of different expressed genes found in analyzed plant species belonging to different land plant clades. Number of genes found in example genomes are shown as red dots. Letters indicate significant differences between groups (P < 0.01, one-way ANOVA, Tukey’s HSD post hoc test). (*C*) GRAS genes per subfamily found in the early diverging land plant genomes from *M. polymorpha, S. fallax*, and *P. patens*, compared to *A. thaliana*. (*D*) Phylogenetic analysis of land plant GRAS proteins using GRAS domains. Support values associated with branches and displayed as bar thickness are maximum likelihood bootstrap values from 1000 replicates.

### Early evolution of DELLA proteins

To elucidate DELLA evolution, we generated a new phylogenetic tree adding previously undetected DELLA proteins from species belonging to different clades across land plant phylogeny (**Fig. 3A**). The N-terminal domain was also excluded from this analysis because the high level of divergence in this region yielded trees that were in conflict with known taxonomic relationships (Supplementary **Fig. 3**). We confirmed the previously reported major DELLA clades corresponding to the eudicot clades RGA and RGL (DELLA1 and DELLA2, respectively), which are fused into a single DELLA1/2 clade in non-eudicot tracheophytes, and the DELLA3 clade, also named DGLLA/SLRL, that is present in all vascular plant lineages. These three clades are found in every major clade analysed, with the sole exception of DELLA1/2 being absent in ferns. This data suggests the occurrence of two main duplication events in *DELLA* genes coinciding with the appearance of tracheophytes and the emergence of eudicots. However, multiple duplications and loses have occurred in specific groups and species, such as the lack of DELLA1 and DELLA3 in *Solanum lycopersicum*.

**Figure 3.**
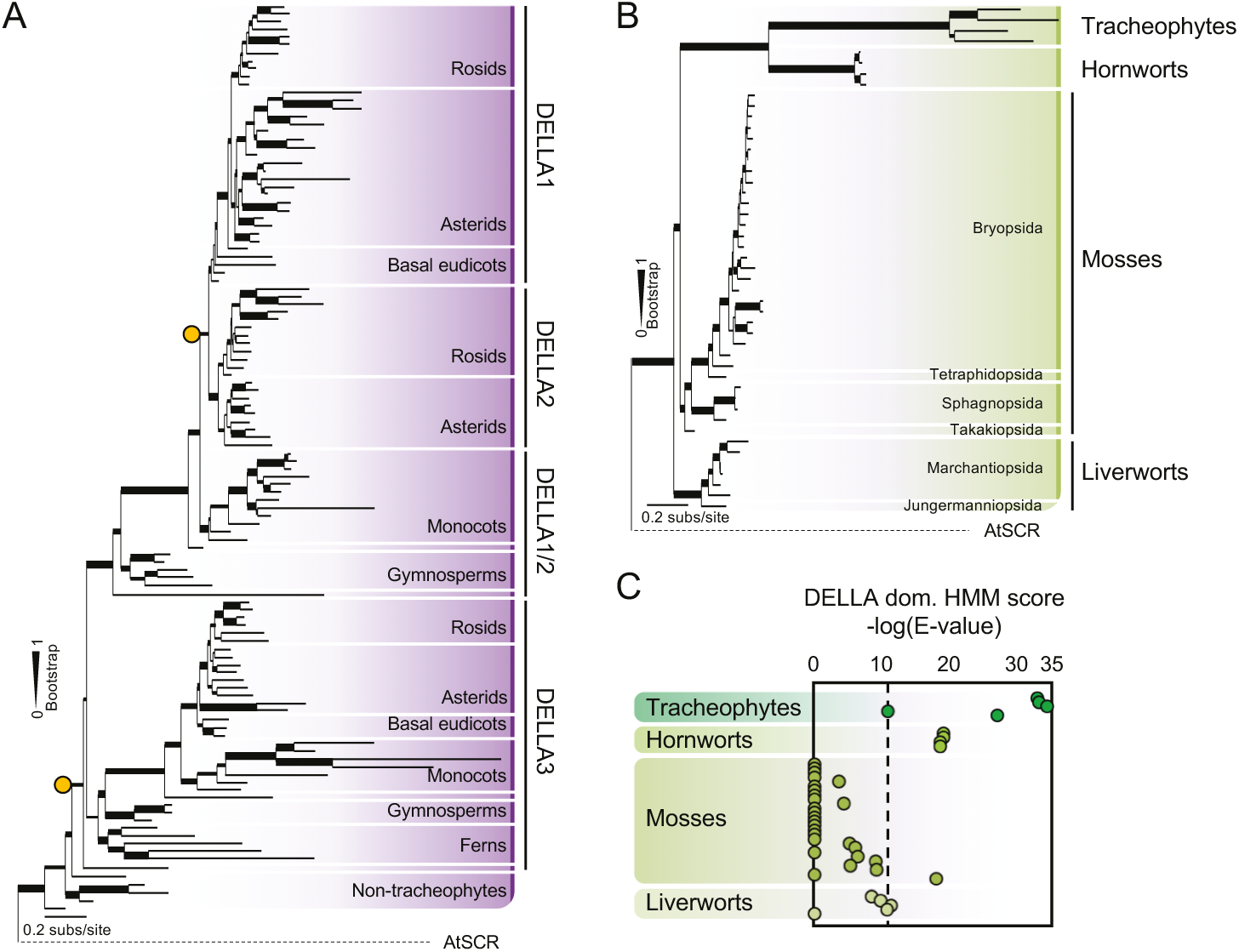
DELLA proteins are present in all extant land plant lineages. (*A*) Phylogenetic analysis of DELLA proteins using GRAS domains of all land plant lineages. Orange circles indicate inferred major duplications within DELLA subfamily. (*B*) Phylogenetic analysis of DELLA proteins using GRAS domains of non-vascular land plant sequences and a few land plant representatives. Support values associated with branches and displayed as bar thickness are maximum likelihood bootstrap values from 1000 replicates. (*C*) Automated search for DELLA motifs in non-vascular land plants using Pfam website. Scores are represented as the negative Log of the E-value retrieved from the search and represented following the phylogenetic position obtained in Fig. 3B. Absence of significant E-values for the presence of Pfam DELLA domain are plotted as 0 in the graph. Dashed line represents the value retrieved for SmDELLA2 DELLA domain. This domain is able to interact with SmGID1a/b while having moderately divergent canonical DELLA domain motifs.

Due to the scarce knowledge of DELLA function in non-vascular land plants, we expanded our search for DELLA sequences in liverworts, mosses and hornworts and analyzed their phylogenetic relationship (**Fig. 3B**). As suggested by the previous tree, no ancient major duplications in DELLAs have been maintained in land plants prior to vasculature emergence. We also detected a type of DELLA sequences that we named DELLA-like proteins (DLP), whose phylogenetic position is unclear and most likely represent a liverwort-specific duplication with no resemblance in their N-terminal domains to those of DELLA proteins (**Supplementary Fig. 3**). Other duplication events have also occurred independently in mosses as in the case of Funariaceae or Sphagnopsida.

### DELLA domain characterization in early diverging land plants

Since the function of DELLA proteins as GA signalling elements requires the presence of specific motifs in their N-terminal domain and it has been proposed that there is no GA pathway in non-vascular plants (Miyazaki et al. 2018), we decided to analyse the occurrence of DELLA motifs in the N-terminal domain of DELLA proteins using automated Pfam HMM domain detection. Contrary to the highly significant scores for the presence of GRAS domains (PF03514; **Supplementary Fig. 3**), the identification of the DELLA Pfam HMM (PF12041) resulted in strongly variable significance values (**Supplementary Table 4**). In tracheophytes, the search was positive with independent E-values usually around 10^-30^, with SmDELLA2 being the higher with an E-value of 1.3x10^-11^ (**Fig. 3C**). SmDELLA2 contains highly divergent DELLA domain motifs but has been shown to be targeted by GID1 upon gibberellin recognition (Hirano et al. 2007), setting an empirical threshold of potentially GID1-targeted DELLA domains. Among non-tracheophyte DELLAs, hornworts scored with E-values of around 10^-19^, indicating that their DELLA domains are very similar to those found in tracheophytes. In agreement with the reported lack of DELLA canonical motifs and functionality of the DELLA-domain in *Physcomitrella patens* (Hirano et al. 2007; Yasumura et al. 2007), mosses show a clear trend toward DELLA Pfam loss, but their earlier diverging moss species contain relatively low E-values, reaching 10^-18^ for *Takakia lepidozioides* DELLA (**Fig. 3C**).

To determine the precise motifs and residues that provide high significance value for the identity of the N-terminal domain, we aligned the corresponding regions of representative sequences from each clade (three liverworts, ten mosses, three hornworts, and three tracheophytes) (**Fig. 4**). The three important motifs for the interaction with the GID1 receptor (and, therefore, for GA signalling) in tracheophytes were differentially conserved among non-vascular plants: liverworts displayed clear DELLA and VHYNPS motifs; most mosses only contain the LEQLE motif, except *T. lepidozioides*, in which only DELLA and VHYNPS are present; and hornwort sequences contain DELLA, LEQLE and VHYNPS motifs.

**Figure 4.**
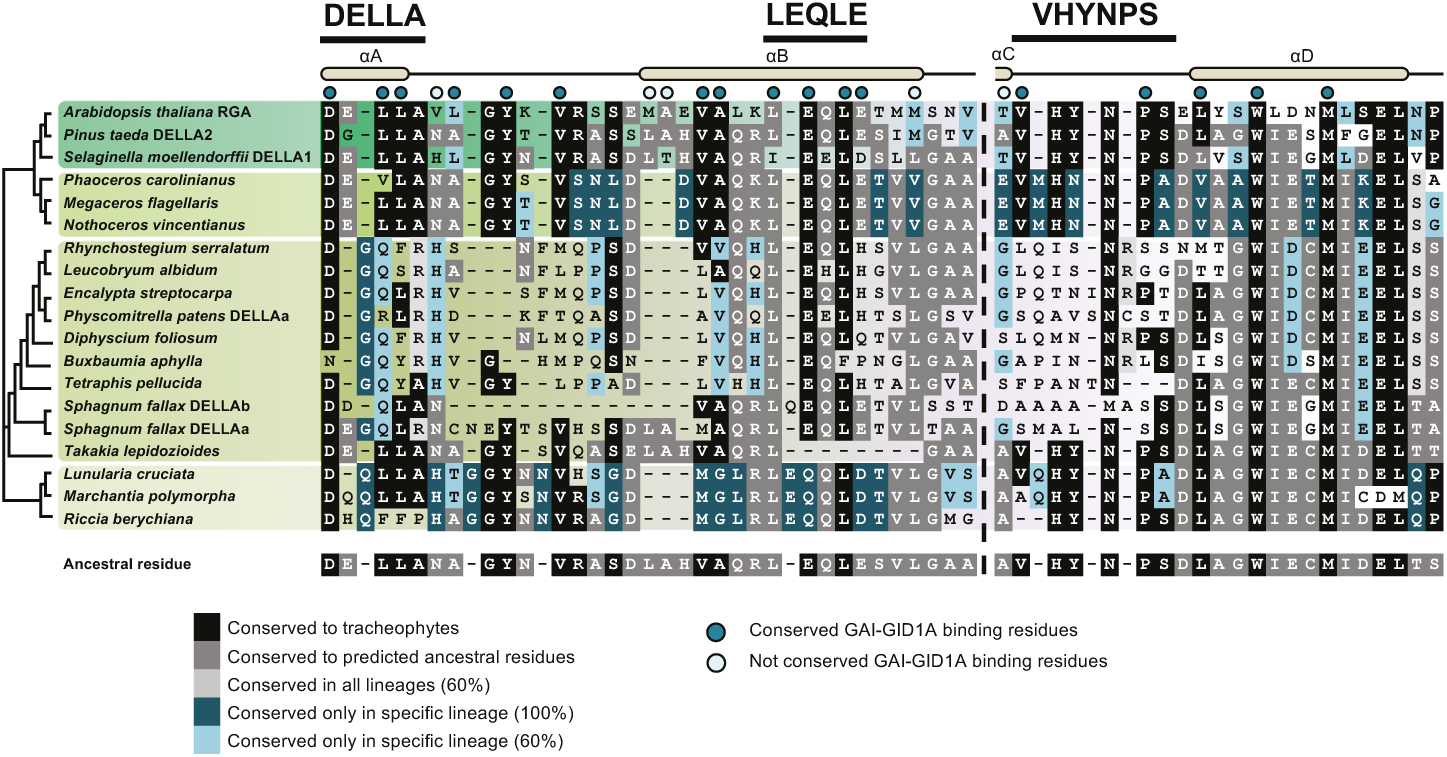
GID1 binding residues in DELLA proteins are highly conserved. Multiple protein sequence alignment showing DELLA amino-terminal region spanning from a helix A to D. In some cases, non-conserved sequences were trimmed to avoid multiple gaps presence. Conservation percentages are based on original alignments. GAI-GID1A binding sites based on Murase et al. 2008. Ancestral DELLA sequence inferred with FastML, only the most probable residues are shown per position.

This distribution of motifs suggests that the N-terminal domain of DELLA proteins was established early in land plant ancestors and maintained during evolution. To confirm this hypothesis, we performed ancestral protein reconstruction by maximum likelihood methods, excluding late divergent moss sequences to avoid bias towards DELLA or VHYNP motif loss. To avoid the lack of consensus in bryophyte relationships and early land plant evolutionary history, we performed the analysis using different phylogenetic relationships among the three early diverging land groups (hornworts, liverworts and mosses), with almost identical results. The predicted ancestral sequences for the land plant common ancestor DELLA N-terminal domain harbours the canonical motifs and is strictly conserved compared to those known in tracheophytes and in hornworts (**Fig. 4)**. Three of the four alpha helices harboured in the N-terminal domain show an ancestral state highly conserved in late diverging DELLA proteins, coinciding with the alpha helices that form the surface interacting with GID1.

Two more pieces of evidence support that the putative function of the ancestral N-terminal domain of DELLA proteins needed to be maintained during evolution: the K_a_/K_s_ ratio is particularly low (around 0.2) precisely in the region that interacts with GID1 in higher plants (**Supplementary Fig. 4A; Supplementary Table 5**); and the characteristic intrinsic disorder of the whole N-terminal domain of DELLA proteins (Sun et al. 2010) is in fact absent in the GID1-interacting region (**Supplementary Fig. 4B,C; Supplementary Table 5**).

### DELLA interaction with GA-signalling components

The solid conservation of the N-terminal domain of DELLA proteins even in land plants that lack the necessary GA signaling elements suggests that the ancestral DELLA was preadapted for subsequent interaction with the GA receptors. To gather additional experimental evidence, we first modelled the structure of putative complexes between AtGID1a and DELLAs from species of different land plant taxa based on the previously described GA_4_-AtGID1a-AtGAI structure. For comparison, the DELLAs were selected from *A. thaliana* (Angiosperma), *S. moellendorffii* (Lycophyta), *Nothoceros vincentianus* (Anthocerotophyta), *P. patens* and *T. lepidozioides* (Bryophyta), and *Marchantia polymorpha* (Marchantiophyta). The structures of the N-terminal domain of the selected DELLAs were modelled and superimposed to the structure of the DELLA-GID1 complex (Kelley et al. 2015). Using this strategy, we were able to determine if the DELLA, LEQLE and VHYNPS motifs of the selected proteins could potentially interact with the AtGID1a protein (**Fig. 5A; Supplementary Fig. 5A**). As expected, the model for *A. thaliana* DELLA (AtRGA) showed a similar interaction with AtGID1a to that observed between AtGAI and AtGID1a. Despite small differences observed on the VHYNP motif of *N. vincentianus* NvDELLA (MHNNP) and the LEQLE motif of *S. moellendorffii* SmDELLA1 (IEELD), both proteins exhibit similar fold and potential interaction with AtGID1a, suggesting that lycophyte and hornwort DELLAs might indeed establish functional interactions with GA receptors. The structure of the SmDELLA1 model in complex with AtGID1 also shows similar interactions except for the LEQLE motif where the latter glutamate (E) is replaced in SmDELLA1 by an aspartate (D) (IEELD). This mutation which should decrease the interaction with K28 (**Fig. 5**) does not seem important because the degradation of SmDELLA1 by GAs has already been described (Hirano et al. 2007; Yasumura et al. 2007). This result can be easily understood by the mobility of the side chain of Lysine (K) 28 which could likely interact with the aspartate residue of SmDELLA1. On the contrary, despite a similar fold, *T. lepidozioides* TlDELLA displayed critical changes in the LEQLE motif (absent in the alignment, substituted by the subsequent LGAAQ sequence in the model) of the αB helix which are predicted to prevent interaction with GID1. In addition, *M. polymorpha* and *P. patens* DELLAs present not only modifications in the αB helix, but also in the DELLA and VHYNP motifs preventing interaction with AtGID1a. In summary, the models indicate that, unlike lycophytes and hornworts, moss and liverwort DELLAs should be unable to interact with AtGID1a.

**Figure 5.**
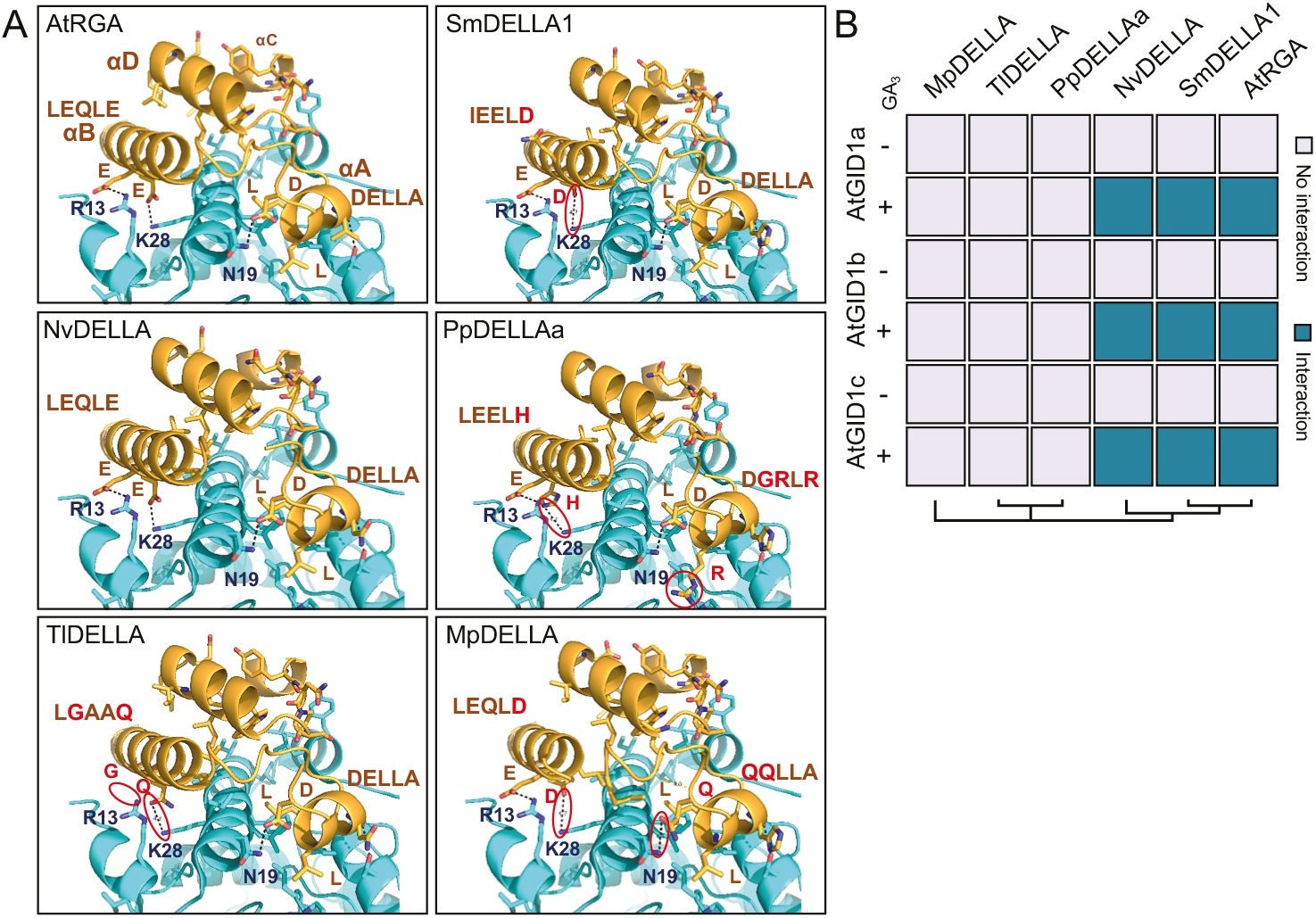
Some non-vascular DELLA proteins can interact with GID1 receptors in a GA dependent manner. (*A*) Predicted structuraI modeI for DELLA-AtGlD1a interaction using AtRGA, SmDELLA1, NvDELLA, PpDELLAa, TIDELLA, and MpDELLA. DELLA structure is shown in yeIIow and AtGlD1a structure in Iight bIue. AtGlD1a residues invoIved in DELLA interaction are written in dark bIue. Residues different to that of AtRGA/GAl in main motifs are presented in red. PossibIe residue to residue interactions affected are pointed with a red circIe (*B*) Yeast-two-hybrid assay resuIts between DELLA proteins and the three Arabidopsis GlD1 receptors with or without GA_3_. Positive interactions are accounted when yeast growth occurs in 5 mM 3-aminotriazoI.

To experimentally test the models, we performed yeast-two-hybrid assays between the six full-length DELLAs and the Arabidopsis GID1 proteins in the presence or absence of GA_3_ (**Fig. 5B; Supplementary Fig. 5B**). As expected, both tracheophytan DELLAs interacted with AtGID1s when GA was present. Interestingly, *N. vincentianus* DELLA was also able to interact with the receptors in a GA-dependent manner. However, moss and liverwort sequences did not show interaction with the receptors either in a GA-dependent or independent way. These results indicate that N-terminal domain conservation is necessary but not sufficient for GID1 interaction. Moreover, the ability of hornwort DELLAs to recognize the Arabidopsis GA receptors in a GA-dependent manner (**Fig. 5B**), and the presence of putative GA receptor sequences in some Anthocerotophyta genomes (**Fig. 1**; **Supplementary Table 2**) suggests that the origin of GA signaling might predate the separation between hornworts and vascular plants.

### An ancestral function of the DELLA N-terminal domain in transcriptional activation

The presence in N-terminal region of the ancestral DELLA protein of structured domains that were necessary for the eventual construction of a GA signaling module begs for an additional function encoded in this region which would explain its conservation in non-vascular plants lacking GA receptors or elaborate GA biosynthesis. Interestingly, although the actual mechanism is still unknown, Arabidopsis DELLAs have been reported to act as transcriptional coactivators in certain developmental contexts (Yoshida et al. 2014; Fukazawa et al. 2014; Marín-de la Rosa et al. 2015). In fact, when we examined the transcriptional status of loci to which AtRGA is bound (Marín-de la Rosa et al. 2015), most of the genes showed a tendency to be induced when DELLAs are active (**Fig, 6A**, **Supplementary Table 6**). It has been reported that expression of full-length rice DELLA fusions to a DNA binding domain in yeast results in the transactivation of the corresponding reporters (Hirano et al. 2012). We have found that this transactivation capacity is conserved in the N-terminal domains of all the DELLAs tested, included those from non-vascular land plants (**Fig. 6B**; **Supplementary Fig. S6A**). Previously, it has been suggested that both DELLA and VHYNP motifs are involved in this activity (Hirano et al. 2012). However, despite the lack of both of these motifs, the N-terminal domain of PpDELLAa is strongly capable of transcriptional activation. The most conserved region among all land plant DELLA N-terminal domains is the αD helix (**Fig. 4**), and we found that deletion of this region (*ΔαD*) in PpDELLAa prevented the induction of reporter expression in yeast, while the αD helix alone was also capable of promoting transactivation in a yeast two-hybrid assay, although this activation is more robust when the whole ordered region (N1) ranging from αA to αC are also present (**Fig. 6C and Supplementary Figure 6B**). To study if this also happens *in planta*, we confirmed these results by performing transient expression assays of a dual luciferase reporter in *Nicotiana benthamiana* (**Fig. 6D, Supplementary Fig. 6C**). The activity of certain transactivation domains from different origins (viral VP16, yeast GAL4 or PHO4, mammalian p53 or NFAT, and others) has been proposed to reside in a particular 9-amino acid transactivation domain (9aaTAD) (Piskacek et al. 2007) which directly interacts with the KIX domain of general transcriptional coactivators like Mediator’s MED15 subunit (Piskacek et al. 2016). Interestingly, the αD helix of the DELLA N-terminal domain displayed a high score in a 9aaTAD evaluation (**Supplementary Fig. 7**).

**Figure 6.**
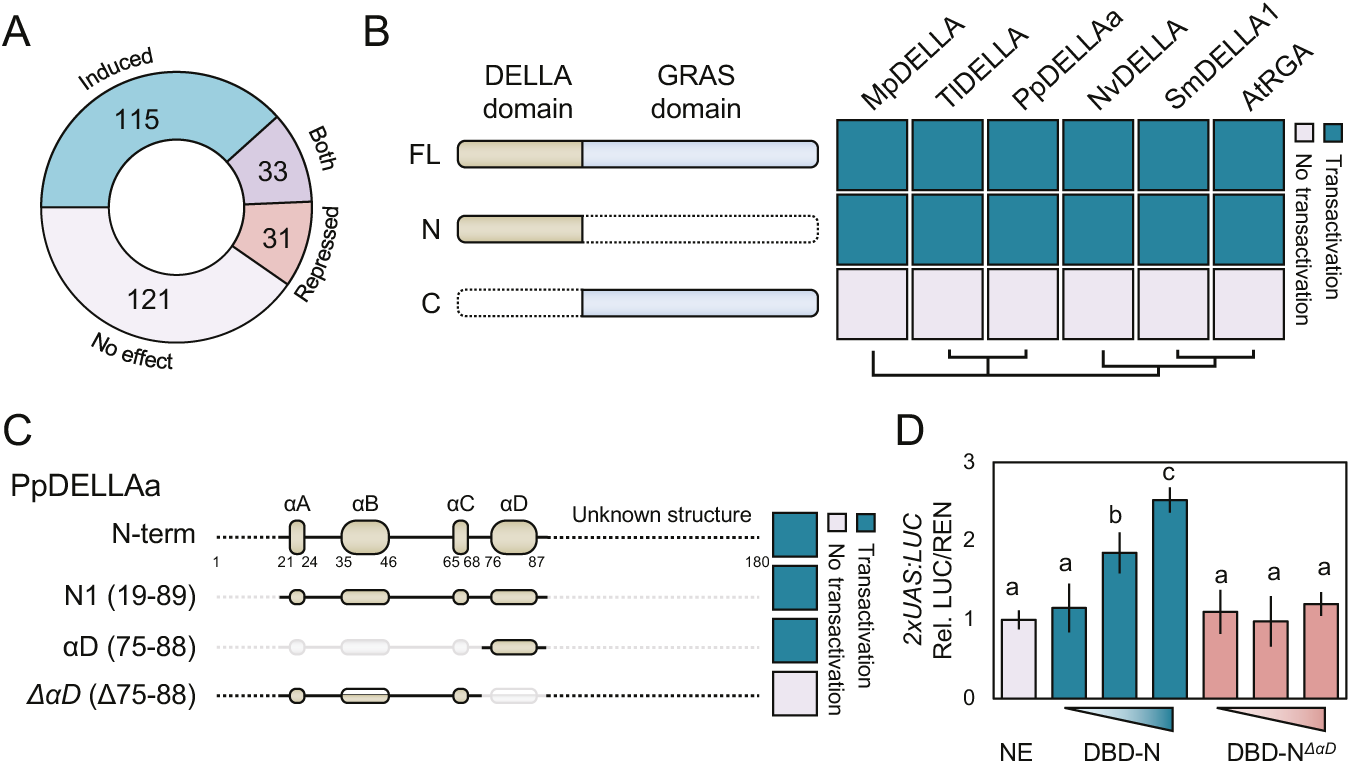
DELLA domain conserved region act as a transcriptional activator domain. (*A*) RGA-bound genes in ChlP-seq assays are enriched in induced versus repressed genes when compared to different transcriptomic data in DELLA induced conditions. ChlP-seq data retrieved from Marfn-de la Rosa et al. 2015; Transcriptomic data obtained from several available datasets. (*B*) Yeast transactivation assay results using DELLA protein full-length coding regions (FL), or truncated versions using either the GRAS domain (C) or the DELLA domain (N). (*C*) Yeast transactivation assay results using different truncated versions of PpDELLAa DELLA domain. Transactivation is accounted when yeast growth occurs in 5 mM 3-aminotriazol. (*D*) Dual luciferase transactivation assays in *Nicotiana benthamiana* leaves using the *LUC* gene under the control of the *Gal* operon UAS promoter as the reporter, and different effector vectors fused to the GAL4 binding domain. NE, no effector; DBD-N, PpDELLAa DELLA domain fused to GAL4 DNA binding domain; DBD-N*ΔlaD*, PpDELLAa Δ75-88 truncated version fused to GAL4 BD. Constitutively expressed *Renilla* luciferase (*REN*) for normalization. Data shown are normalized to NE value and represent the average of three biological replicates. Error bars represent standard deviation. Letters indicate significant differences between groups (P < 0.01, one-way ANOVA, Tukey’s HSD post hoc test).

## Discussion

The work presented here provides new clues about the origin of DELLA proteins in the common ancestor of all land plants, and a possible mechanism by which these proteins became GA signaling elements in vascular plants, after the emergence of the GID1 GA receptor.

Previously, putative DELLA proteins had been reported in at least two non-vascular plant species: the moss *P. patens*, and the liverwort *M. polymorpha* (Hirano et al. 2007; Yasumura et al. 2007; Bowman et al. 2017). Our extensive phylogenetic analyses have not only confirmed the widespread presence of clearly defined DELLA proteins in all clades of non-vascular plants including hornworts, but also add two important pieces of new information: (i) since all land plant GRAS proteins are monophyletic, the origin of DELLA proteins unequivocally coincides with the colonization of land by plants; and (ii) the N-terminal domain is conserved in the vast majority of DELLAs, including those in non-vascular plants. This observation contradicts the previous assumption that the recruitment of DELLAs to GA signaling was due to the appearance of GID1-interacting motifs in this N-terminal region in vascular plants (Hirano et al. 2007; Yasumura et al. 2007). This assumption was largely based on the absence of the ‘DELLA’ and ‘VHYNPS’ motifs in PpDELLA; but the presence of these important motifs in a basal moss species, like *T. lepidozioides,* in all the hornwort DELLA sequences analyzed, and in the reconstructed ancestral DELLA protein sequence, suggest a most likely scenario in which the ancestral DELLA contained most of the motifs that would later be useful to establish the interaction with the GID1 receptor.

The establishment of the GID1-DELLA interaction constitutes, as previously reasoned (Sun 2011), the key event that connects the ancestral DELLA activity with the newly emerging GA signaling. In this respect, our work contributes with the finding of GID1-like sequences in hornworts that are phylogenetically close to bona-fide GID1 receptors, and the observation that the N-terminal domains of hornwort DELLAs display the intrinsic ability to interact with a vascular plant GID1 receptor in a GA-dependent manner. Therefore, at least two possible models can be contemplated: either GA signaling emerged in a common ancestor of hornworts and vascular plants, or it emerged independently in vascular plants and in hornworts, and the similar behaviour is caused by functional convergence. A third possibility would be that a putative GA-independent GID1-DELLA module in the ancestor of all land plants would have been lost in different clades, but this is highly unlikely based on recent evidence about GID1 evolution(Yoshida et al. 2018). The origin of the participation of DELLAs in GA signaling requires a more complete picture. Future work is needed to answer several key questions: (i) do hornwort GID1-like proteins behave as GA receptors?; (ii) can hornwort DELLAs interact with hornwort GID1-like proteins in a GA-dependent manner (or is any other hormone-like molecule perceived by GID1-like proteins)?; (iii) is DELLA activity regulated in non-vascular plants by other GA-related compounds which are considered as GA precursors in vascular plants? Curiously, 3-hydroxy-kaurenoic acid has been proposed as a plant growth regulator in *P. patens*(Miyazaki et al. 2018), indicating a functional role for at least this metabolite in GA metabolism, but it is currently unknown if this function is conserved in other non-vascular plants.

Our results suggest that the main driving force for the conservation of the N-terminal regions of DELLA proteins has been its role in transcriptional activation, and its eventual co-option by the GID1 receptor allowed hormonal regulation of DELLA stability. Although molecular exploitation has been described as an evolutionary strategy to expand hormone-receptor complexity (Bridgham et al. 2006; Baker et al. 2013), the function of the evolving receptor was equivalent to the ancestral one (i.e., interaction with the ligand). However, the origin of the GA signaling pathway illustrates how molecular exploitation can occur upon domains with completely unrelated functions. Curiously, the co-existence of degron and transactivation motifs in the same stretches of residues has been described in several mammalian transcriptional activators (Salghetti 2000; Salghetti et al. 2001). In contrast, this degron-TAD overlap has not been studied in plants but some examples can be identified, such as MYC2, in which a degron is found within the MID domain, the transactivation domain (Zhai et al. 2013). Therefore, DELLAs would have these two functions encoded in a single region, and interaction with GID1 has been reported not only to promote DELLA degradation, but also prevent transactivation by DELLAs (Hirano et al. 2012). In summary, the coincidence of transactivation and protein stability regulation in a single protein domain is a widespread property and has independently emerged several times during evolution and through different molecular mechanisms.

## Materials and Methods

### Identification of GRAS and GA signalling elements sequences in plants

GRAS homolog sequences were searched in Phytozome, OneKP, and specific databases for the charopyte *Klebsormidium flaccidum* (reassigned as *K. nitens*), red algae, and the glaucophyte *Cyanophora paradoxa* (Supplementary Table 7). *A. thaliana* and *P. patens* previously identified GRAS sequences were re-checked and used as query in a BLASTP initial search. In short, proteomes were examined using a BLASTP local blast search using an e-value cut-off of 0.1 in most cases, further raised to 10 in red algae, chlorophytes o *C. paradoxa* in order to avoid missing highly diverging GRAS sequences. Initial results were first subjected to reciprocal BLAST. Subsequently, the results were manually checked using SMART (http://smart.emblheidelberg.de/) and Pfam (http://pfam.sanger.ac.uk/search) to ensure GRAS domain presence. For GID1 and GID2 related sequences, Phytozome and OneKP databases were analysed as mentioned, using *A. thaliana* and *S. moellendorffii* previously identified protein sequences as query. In this case, only reciprocal BLAST was used. Manual curation of incomplete annotations was performed when needed using either transcriptomic data from the same or the closest species orthologs when available.

### Phylogenetic analysis

Sequences were aligned using the MUSCLE algorithm (Edgar 2004) included in the SeaView GUI (Gouy et al. 2010), with 16 iterations, default clustering methods, gap open score of - 2.7, and hydrophobicity multiplier of 1.2, followed by manual curation. For phylogenetic reconstruction, C-terminal GRAS domains were used, and ambiguously aligned regions manually trimmed. In the case of DELLA phylogenetic analysis, AtSCR was included in the final alignments before tree reconstruction using MAFFT v7 method L-INS-i (Katoh and Standley 2013). ProtTest v3.4.2 (Darriba et al. 2011) was used on final multiple sequence alignments to select best-fit models of amino acid replacement using the AIC model for ranking. Maximum likelihood tree in Fig. 2a was produced with RAxML 8.2.3 using the LG PROTGAMMA model (Stamatakis 2014). The rest of ML trees were produced with PhyML v3.1 (Guindon et al. 2010), using the best scored model of amino acid substitution. Statistical significance was evaluated by bootstrap analysis of 1000 replicates in all cases with the sole exception of Supplemental Figure 1, which was evaluated by SH-like approximate likelihood ratio test (aLRT). Phylogenetic tree graphical representations were initially generated using FigTree (version 1.4.3) software (http://tree.bio.ed.ac.uk/software/figtree/), and final cartoons edited manually.

Original sequences, raw and trimmed alignments, and trees are available at https://data.mendeley.com/datasets/bjcp6ggjk9/draft?a=6c3474c7-6b37-47d0-ad3d-19ad9c062627

### Ancestral sequence reconstruction

The ancestral state for each codon position in the DELLA N-terminal domain was determined using MEGA7 (Kumar et al. 2015). Nucleotide coding sequences aligned following the previous result of the corresponding aminoacids alignment. Ancestral sequence inference was then performed using Maximum Likelihood, including four different predefined tree topologies around non vascular plants: (i) monophyletic bryophyta, (ii) a moss–liverwort sister clade to other embryophytes; (iii) liverwort–moss sister clade to tracheophytes; and (iv) hornworts, mosses, and liverworts as successive sister lineages to tracheophytes(Puttick et al. 2018). Finally, the Tamura-Nei model of nucleotide substitution was used for ancestral state inference. Gap residues were trimmed if absent in more than 90% of the sequences.

### Codon selection and protein disorder analysis

Analysis of selection was performed using the web-based interface Selecton v2.2 (Stern et al. 2007). M8 and M8a models of selection were used to calculate the ratio between the rates of non-synonymous (K_a_) and synonymous substitution (K_s_) in previously constructed codon-based nucleotide alignments. Likelihood scores estimated by the models were evaluated by log-likelihood ratio testing with degree of freedom (df)=1, followed by Bayesian prediction of undergoing positive approach. Prediction of disorder per residue was performed with the ANCHOR web tool (https://iupred2a.elte.hu/) (Dosztányi et al. 2009). Mean predicted disorder values per residue were calculated based on the back-translated codon-based alignment using AtRGA, SmDELLA1, NvDELLA, PpDELLAa, TlDELLA, and MpDELLA. Raw data is included in **Supplemental Table 5**.

### Protein structure prediction

The N-terminal regions of all DELLA proteins were modelled with 100% confidence using AtGAI (PDB code 2ZSH) (Murase et al. 2008) as template using the PHYRE2 program (Kelley et al. 2015) and visualized with PyMOL software (The PyMOL Molecular Graphics System, Version 2.0 Schrödinger, LLC.).

### Transactivation domain prediction

AtRGA, SmDELLA1, NvDELLA, PpDELLAa, TlDELLA, MpDELLA protein sequences were analysed with a nine-amino-acid transactivation domain (9aa TAD) prediction tool (http://www.med.muni.cz/9aaTAD/index.php) (Piskacek et al. 2007), using the “less stringent” pattern. Cumulative probabilities of 9aaTAD for all the proteins were plotted versus an amino acid alignment of the DELLA domain.

### Yeast-two-hybrid assay

Arabidopsis GID1 was fused to the Gal4-DNA Binding Domain (DBD) in the pGBKT7-GW vector as bait, and DELLA full-length ORFs from the different species were fused to the Gal4-Activation Domain (AD) in pGADT7-GW. DELLAs and GID1 ORFs were either amplified by PCR using sequence-specific primers (**Supplementary Table 9**) or synthesized as gBlocks^®^ (I.D.T) and transferred to pDONR221 or pDONR207 via BP Clonase II (Invitrogen), or to pCR8 via TOPO^®^TA cloning^®^ (Invitrogen) to create entry vectors. Final constructs were made by recombining entry clones to GATEWAY destination vectors via LR Clonase II (Invitrogen). Direct interaction assays in yeast were performed following Clontech’s small-scale yeast transformation procedure. Strain Y187 was transformed with pGADT7 derived expression vectors, while strain Y2HGold was transformed with pGBKT7 vectors, and selected in SD medium without Leu or Trp, respectively. Subsequently, diploid cells were obtained by mating and selection in SD medium lacking both Leu and Trp. Interaction tests were done in SD medium lacking Leu, Trp and His, in the presence of different concentrations of 3-aminotriazol (3-AT) (Sigma-Aldrich). To assess GA dependent interaction the medium was supplemented (or not) with 100 µM GA_3_.

### Yeast transactivation assay

DELLA ORFs were obtained as described above. DELLA N-end clones and C-end clones were obtained by PCR amplification using sequence-specific primers (**Supplementary Table 9**) and transferred to pDONR207 via BP Clonase II (Invitrogen). Entry clones were then used to create Gal4-DBD fusions in the pGBKT7-GW vector via LR Clonase II (Invitrogen), which was transformed into yeast strain Y2HGold and transformants were selected in SD medium lacking Trp. Transactivation tests were performed in SD medium without Trp and His, and increasing 3-AT concentrations as indicated.

### *In planta* transient transactivation assay

A reporter construct containing 2xGal4 UAS followed by a 35S minimal promoter (Gendron et al. 2012) was amplified using sequence-specific primers (**Supplementary Table 9**) and cloned upstream of the firefly luciferase gene (LUC) in pGreenII 0800-LUC (Hellens et al. 2005). The effector vectors were obtained by amplifying the GAL4 DBD-DELLA N-end fusions generated in pGBKT7 vectors as a unique PCR product with proper restriction site overhangs and ligated into pFGC5941 (http://www.ChromDB.org), between *Xho*I and *Spe*I. The GAL4 DBD control construct was obtained by excising the RGA-N fragment from the DBD-RGA-N vector. Transient expression in *Nicotiana benthamiana* leaves was carried as previously reported (Marín-de la Rosa et al. 2015). Firefly and the control *Renilla* luciferase activities were assayed in extracts from 1-cm in diameter leaf discs, using the Dual-Glo Luciferase Assay System (Promega) and quantified in a GloMax 96 Microplate Luminometer (Promega).

## Supporting information

## Acknowledgments

We thank the members of the Hormone Signaling and Plasticity Lab at IBMCP (http://plasticity.ibmcp.csic.es/) for useful discussions and suggestions. Work in our laboratories was supported by grants BFU2016-80621-P of the Spanish Ministry of Economy, Industry and Competitiveness, and H2020-MSCA-RISE-2014-644435 of the European Union. JH-G and AB-M hold Fellowships of the Spanish Ministry of Education, Culture and Sport FPU15/01756 and FPU14/01941, respectively.

## References

Baker ME, Funder JW, Kattoula SR. 2013. Evolution of hormone selectivity in glucocorticoid and mineralocorticoid receptors. J Steroid Biochem Mol Biol. 137:57–70. doi:10.1016/j.jsbmb.2013.07.009.

Bowman JL, Kohchi T, Yamato KT, Jenkins J, Shu S, Ishizaki K, Yamaoka S, Nishihama R, Nakamura Y, Berger F, et al. 2017. Insights into Land Plant Evolution Garnered from the Marchantia polymorpha Genome. Cell. 171(2):287–304.e15. doi:10.1016/j.cell.2017.09.030.

Bridgham JT, Carroll SM, Thornton JW. 2006. Evolution of hormone-receptor complexity by molecular exploitation. Science (80-). 312(5770):97–101. doi:10.1126/science.1123348.

Briones-Moreno A, Hernández-García J, Vargas-Chávez C, Romero-Campero FJ, Romero JM, Valverde F, Blázquez MA. 2017. Evolutionary Analysis of DELLA-Associated Transcriptional Networks. Front Plant Sci. 8. doi:10.3389/fpls.2017.00626.

Darriba D, Taboada GL, Doallo R, Posada D. 2011. ProtTest 3: fast selection of best-fit models of protein evolution. Bioinformatics. 27(8):1164–5. doi:10.1093/bioinformatics/btr088.

Delaux P-M, Radhakrishnan G V., Jayaraman D, Cheema J, Malbreil M, Volkening JD, Sekimoto H, Nishiyama T, Melkonian M, Pokorny L, et al. 2015. Algal ancestor of land plants was preadapted for symbiosis. Proc Natl Acad Sci. 112(43):13390–13395. doi:10.1073/pnas.1515426112.

Dill A. 2004. The Arabidopsis F-Box Protein SLEEPY1 Targets Gibberellin Signaling Repressors for Gibberellin-Induced Degradation. PLANT CELL ONLINE. 16(6):1392–1405. doi:10.1105/tpc.020958.

Dill A, Jung HS, Sun TP. 2001. The DELLA motif is essential for gibberellin-induced degradation of RGA. Proc Natl Acad Sci U S A. 98(24):14162–l14167. doi:10.1073/pnas.251534098.

Dosztányi Z, Mészáros B, Simon I. 2009. ANCHOR: web server for predicting protein binding regions in disordered proteins. Bioinformatics. doi:10.1093/bioinformatics/btp518.

Edgar RC. 2004. MUSCLE: multiple sequence alignment with high accuracy and high throughput. Nucleic Acids Res. 32(5):1792–1797. doi:10.1093/nar/gkh340. [accessed 2013 Mar 6]. http://www.pubmedcentral.nih.gov/articlerender.fcgi?artid=390337&tool=pmcentrez&rendertype=abstract.

Engstrom EM. 2011. Phylogenetic analysis of GRAS proteins from moss, lycophyte and vascular plant lineages reveals that GRAS genes arose and underwent substantial diversification in the ancestral lineage common to bryophytes and vascular plants. Plant Signal Behav. 6(6):850–854. doi:10.4161/psb.6.6.15203.

Floss DS, Levy JG, Levesque-Tremblay V, Pumplin N, Harrison MJ. 2013. DELLA proteins regulate arbuscule formation in arbuscular mycorrhizal symbiosis. Proc Natl Acad Sci. 110(51):E5025–E5034. doi:10.1073/pnas.1308973110.

Fukazawa J, Teramura H, Murakoshi S, Nasuno K, Nishida N, Ito T, Yoshida M, Kamiya Y, Yamaguchi S, Takahashi Y. 2014. DELLAs Function as Coactivators of GAI-ASSOCIATED FACTOR1 in Regulation of Gibberellin Homeostasis and Signaling in *Arabidopsis*. Plant Cell. doi:10.1105/tpc.114.125690.

Gendron JM, Pruneda-Paz JL, Doherty CJ, Gross AM, Kang SE, Kay S a. 2012. Arabidopsis circadian clock protein, TOC1, is a DNA-binding transcription factor. Proc Natl Acad Sci U S A. 109(8):3167–72. doi:10.1073/pnas.1200355109. [accessed 2014 May 8]. http://www.pubmedcentral.nih.gov/articlerender.fcgi?artid=3286946&tool=pmcentrez&rendertype=abstract.

Gomi K, Sasaki A, Itoh H, Ueguchi-Tanaka M, Ashikari M, Kitano H, Matsuoka M. 2004. GID2, an F-box subunit of the SCF E3 complex, specifically interacts with phosphorylated SLR1 protein and regulates the gibberellin-dependent degradation of SLR1 in rice. Plant J. 37(4):626–634. doi:10.1111/j.1365-313X.2003.01990.x.

Gouy M, Guindon S, Gascuel O. 2010. SeaView Version 4: A Multiplatform Graphical User Interface for Sequence Alignment and Phylogenetic Tree Building. Mol Biol Evol. 27(2):221–224. doi:10.1093/molbev/msp259.

Griffiths J, Murase K, Rieu I, Zentella R, Zhang Z-L, Powers SJ, Gong F, Phillips AL, Hedden P, Sun T, et al. 2006. Genetic characterization and functional analysis of the GID1 gibberellin receptors in Arabidopsis. Plant Cell. 18:3399–3414. doi:10.1105/tpc.106.047415.

Guindon S, Gascuel O, Dufayard J-F, Lefort V, Anisimova M, Hordijk W. 2010. New Algorithms and Methods to Estimate Maximim-Likelihood Phylogenies: Assessing the Performance of PhyML 3.0. Syst Biol. 59(3):307–321. doi:10.1093/sysbio/syq010.

Hayashi K -i., Horie K, Hiwatashi Y, Kawaide H, Yamaguchi S, Hanada A, Nakashima T, Nakajima M, Mander LN, Yamane H, et al. 2010. Endogenous Diterpenes Derived from ent-Kaurene, a Common Gibberellin Precursor, Regulate Protonema Differentiation of the Moss Physcomitrella patens. PLANT Physiol. 153(3):1085–1097. doi:10.1104/pp.110.157909.

Hellens RP, Allan AC, Friel EN, Bolitho K, Grafton K, Templeton MD, Karunairetnam S, Gleave AP, Laing W a. 2005. Transient expression vectors for functional genomics, quantification of promoter activity and RNA silencing in plants. Plant Methods. 1:13. doi:10.1186/1746-4811-1-13. [accessed 2014 May 8]. http://www.pubmedcentral.nih.gov/articlerender.fcgi?artid=1334188&tool=pmcentrez&rendertype=abstract.

Hirano K, Kouketu E, Katoh H, Aya K, Ueguchi-Tanaka M, Matsuoka M. 2012. The suppressive function of the rice della protein SLR1 is dependent on its transcriptional activation activity. Plant J. 71(3):443–453. doi:10.1111/j.1365-313X.2012.05000.x.

Hirano K, Nakajima M, Asano K, Nishiyama T, Sakakibara H, Kojima M, Katoh E, Xiang H, Tanahashi T, Hasebe M, et al. 2007. The GID1-mediated gibberellin perception mechanism is conserved in the Lycophyte Selaginella moellendorffii but not in the Bryophyte Physcomitrella patens. Plant Cell. 19(10):3058–3079. doi:10.1105/tpc.107.051524.

Itoh H, Ueguchi-Tanaka M, Sato Y, Ashikari M, Matsuoka M. 2002. The gibberellin signaling pathway is regulated by the appearance and disappearance of SLENDER RICE1 in nuclei. Plant Cell. 14(January):57–70. doi:10.1105/tpc.010319.

Katoh K, Standley DM. 2013. MAFFT multiple sequence alignment software version 7: Improvements in performance and usability. Mol Biol Evol. doi:10.1093/molbev/mst010.

Kelley LA, Mezulis S, Yates CM, Wass MN, Sternberg MJE. 2015. The Phyre2 web portal for protein modeling, prediction and analysis. Nat Protoc. doi:10.1038/nprot.2015.053.

Kumar S, Stecher G, Tamura K, Stecher G, Peterson D, Filipski A, Kumar S. 2015. MEGA7: Molecular Evolutionary Genetics Analysis version 7.0. Mol Biol Evol. submitted(12):2725–2729. doi:10.1093/molbev/mst197.

Locascio A, Blázquez MA, Alabadí D. 2013. Genomic analysis of DELLA protein activity. Plant Cell Physiol. 54(8):1229–37. doi:10.1093/pcp/pct082. [accessed 2014 Dec 23]. http://www.ncbi.nlm.nih.gov/pubmed/23784221

Marín-de la Rosa N, Pfeiffer A, Hill K, Locascio A, Bhalerao RP, Miskolczi P, Grønlund AL, Wanchoo-Kohli A, Thomas SG, Bennett MJ, et al. 2015. Genome Wide Binding Site Analysis Reveals Transcriptional Coactivation of Cytokinin-Responsive Genes by DELLA Proteins. Yu H, editor. PLoS Genet. 11(7):e1005337. doi:10.1371/journal.pgen.1005337.

Marín-de la Rosa N, Sotillo B, Miskolczi P, Gibbs DJ, Vicente J, Carbonero P, Onate-Sanchez L, Holdsworth MJ, Bhalerao R, Alabadi D, et al. 2014. Large-Scale Identification of Gibberellin-Related Transcription Factors Defines Group VII ETHYLENE RESPONSE FACTORS as Functional DELLA Partners. Plant Physiol. 166(2):1022–1032. doi:10.1104/pp.114.244723.

Mcginnis KM, Thomas SG, Soule JD, Strader LC, Zale JM, Sun T, Steber CM, Sly GG. 2003. The Arabidopsis *SLEEPY1* gene encodes a putative F-box subunit of an SCF E3 ubiquitin ligase. Plant Cell. 15(May):1120–1130. doi:10.1105/tpc.010827.increased.

Miyazaki S, Hara M, Ito S, Tanaka K, Asami T, Hayashi K, Kawaide H, Nakajima M. 2018. An Ancestral Gibberellin in a Moss Physcomitrella patens. Mol Plant. 11(8):1097–1100. doi:10.1016/j.molp.2018.03.010.

Murase K, Hirano Y, Sun T, Hakoshima T. 2008. Gibberellin-induced DELLA recognition by the gibberellin receptor GID1. Nature. 456:459–463. doi:10.1038/nature07519.

Peng J, Harberd NP. 1997. Gibberellin deficiency and response mutations suppress the stem elongation phenotype of phytochrome-deficient mutants of Arabidopsis. Plant Physiol. 113(4):1051–1058. doi:10.1104/pp.113.4.1051. [accessed 2015 Jan 4]. http://www.pubmedcentral.nih.gov/articlerender.fcgi?artid=158228&tool=pmcentrez&rendertype=abstract.

Peng J, Richards DE, Hartley NM, Murphy GP, Devos KM, Flintham JE, Beales J, Fish LJ, Worland AJ, Pelica F, et al. 1999. ‘Green revolution’ genes encode mutant gibberellins response modulators. Nature. 400:256–261. doi:10.1038/22307.

Piskacek M, Havelka M, Rezacova M, Knight A. 2016. The 9aaTAD transactivation domains: From Gal4 to p53. PLoS One. 11(9). doi:10.1371/journal.pone.0162842.

Piskacek S, Gregor M, Nemethova M, Grabner M, Kovarik P, Piskacek M. 2007. Nine-amino-acid transactivation domain: Establishment and prediction utilities. Genomics. 89(6):756–768. doi:10.1016/j.ygeno.2007.02.003.

Puttick MN, Morris JL, Williams TA, Cox CJ, Edwards D, Kenrick P, Pressel S, Wellman CH, Schneider H, Pisani D, et al. 2018. The Interrelationships of Land Plants and the Nature of the Ancestral Embryophyte. Curr Biol. doi:10.1016/j.cub.2018.01.063.

Richards DE, Peng J, Harberd NP. 2000. Plant GRAS and metazoan STATs: One family? BioEssays. 22(6):573–577. doi:10.1002/(SICI)1521-1878(200006)22:6573::AID-BIES10>3.0.CO;2-H.

Salghetti SE. 2000. Functional overlap of sequences that activate transcription and signal ubiquitin-mediated proteolysis. Proc Natl Acad Sci. 97(7):3118–3123. doi:10.1073/pnas.050007597.

Salghetti SE, Caudy AA, Chenoweth JG, Tansey WP. 2001. Regulation of transcriptional activation domain function by ubiquitin. Science (80-). 293(5535):1651–1653. doi:10.1126/science.1062079.

Sasaki A, Itoh H, Gomi K, Ueguchi-Tanaka M, Ishiyama K, Kobayashi M, Jeong DH, An G, Kitano H, Ashikari M, et al. 2003. Accumulation of phosphorylated repressor for Gibberellin signaling in an F-box mutant. Science (80-). doi:10.1126/science.1081077.

Stamatakis A. 2014. RAxML version 8: A tool for phylogenetic analysis and post-analysis of large phylogenies. Bioinformatics. 30(9):1312–1313. doi:10.1093/bioinformatics/btu033.

Stern A, Doron-Faigenboim A, Erez E, Martz E, Bacharach E, Pupko T. 2007. Selecton 2007: Advanced models for detecting positive and purifying selection using a Bayesian inference approach. Nucleic Acids Res. 35(SUPPL.2). doi:10.1093/nar/gkm382.

Sun TP. 2011. The molecular mechanism and evolution of the GA-GID1-DELLA signaling module in plants. Curr Biol. 21(9):R338–R345. doi:10.1016/j.cub.2011.02.036.

Sun X, Jones WT, Harvey D, Edwards PJB, Pascal SM, Kirk C, Considine T, Sheerin DJ, Rakonjac J, Oldfield CJ, et al. 2010. N-terminal domains of DELLA proteins are intrinsically unstructured in the absence of interaction with GID1/gibberellic acid receptors. J Biol Chem. 285(15):11557–11571. doi:10.1074/jbc.M109.027011.

Tanaka J, Yano K, Aya K, Hirano K, Takehara S, Koketsu E, Ordonio RL, Park S-H, Nakajima M, Ueguchi-Tanaka M, et al. 2014. Antheridiogen determines sex in ferns via a spatiotemporally split gibberellin synthesis pathway. Science (80-). 346(6208):469–473. doi:10.1126/science.1259923.

Ueguchi-Tanaka M, Ashikari M, Nakajima M, Itoh H, Katoh E, Kobayashi M, Chow T, Hsing YC, Kitano H, Yamaguchi I, et al. 2005. GIBBERELLIN INSENSITIVE DWARF1 encodes a soluble receptor for gibberellin. Nature. 437:693–698. doi:10.1038/nature04028.

Vera-Sirera F, Gomez MD, Perez-Amador MA. 2015. DELLA Proteins, a Group of GRAS Transcription Regulators that Mediate Gibberellin Signaling. In: Plant Transcription Factors. Elsevier. p. 313–328. [accessed 2015 Sep 15]. http://www.sciencedirect.com/science/article/pii/B9780128008546000208.

Yasumura Y, Crumpton-Taylor M, Fuentes S, Harberd NP. 2007. Step-by-Step Acquisition of the Gibberellin-DELLA Growth-Regulatory Mechanism during Land-Plant Evolution. Curr Biol. 17(14):1225–1230. doi:10.1016/j.cub.2007.06.037.

Yoshida H, Hirano K, Sato T, Mitsuda N, Nomoto M, Maeo K, Koketsu E, Mitani R, Kawamura M, Ishiguro S, et al. 2014. DELLA protein functions as a transcriptional activator through the DNA binding of the INDETERMINATE DOMAIN family proteins. Proc Natl Acad Sci1. 111(21):7861–7866. doi:10.1073/pnas.1321669111.

Yoshida H, Tanimoto E, Hirai T, Miyanoiri Y, Mitani R, Kawamura M, Takeda M, Takehara S, Hirano K, Kainosho M, et al. 2018. Evolution and diversification of the plant gibberellin receptor GID1. Proc Natl Acad Sci. doi:10.1073/pnas.1806040115.

Zhai Q, Yan L, Tan D, Chen R, Sun J, Gao L, Dong MQ, Wang Y, Li C. 2013. Phosphorylation-Coupled Proteolysis of the Transcription Factor MYC2 Is Important for Jasmonate-Signaled Plant Immunity. PLoS Genet. 9(4). doi:10.1371/journal.pgen.1003422.

Zhang D, Iyer LM, Aravind L. 2012. Bacterial GRAS domain proteins throw new light on gibberellic acid response mechanisms. Bioinformatics. 28(19):2407–11. doi:10.1093/bioinformatics/bts464. [accessed 2015 Sep 9]

