## Supplementary material for "Origin of gibberellin-dependent transcriptional regulation by molecular exploitation of a transactivation domain in DELLA proteins"

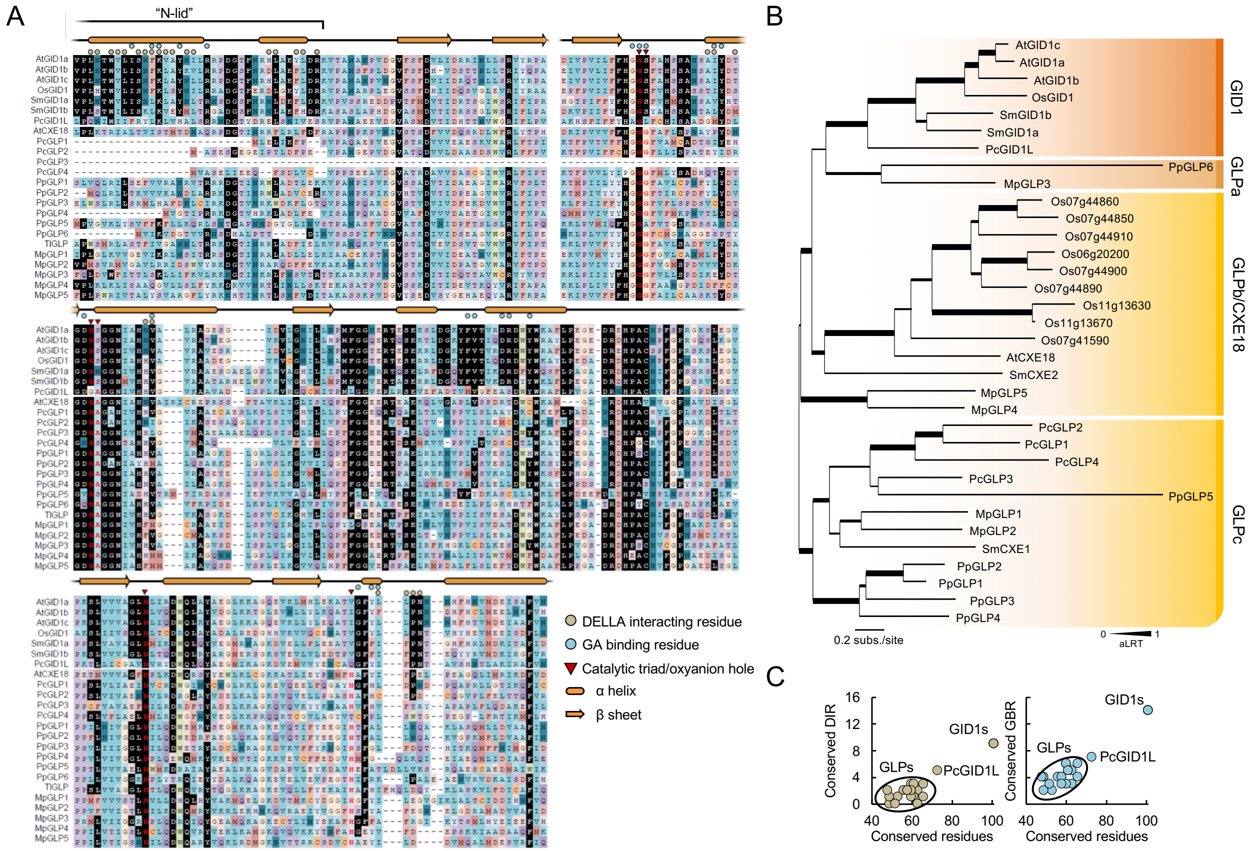


**Supplemental Figure 1. GID1 *bona fide* orthologs may be unique to vascular plants**. (*A*) Multiple sequence alignment of GID1 and GID1-like proteins from several land plant species. GID1 structure and DELLA/GA binding residues based on Murase et al. 2008. Black background denotes highly conserved residues in vascular plants. Red letters indicate catalytic triad and oxyanion hole residues from the α/β hydrolase family. (*B*) Phylogenetic analysis of GID1 proteins. Support values associated with branches and displayed as bar thickness are SH-like approximate likelihood ratio test scores (aLRT). Unlike other non-vascular plants, the hornwort *Phaeoceros carolinianus* harbours a sequence which aligns in the same clade as bona-fide GID1 GA receptors, and contains mutations in the catalytic triad that resemble those of GA receptors (as shown in A). We have named it PcGID1L. (*C*) Comparison between the number of DELLA interacting residue (DIR) or GA binding residue (GBR) conservation and the total number of residues conserved compared to vascular plant GID1s. Only strictly conserved residues in vascular plants are counted (black background in Sup. Fig. 1A). GLPs from non-vascular land plants are encircled.

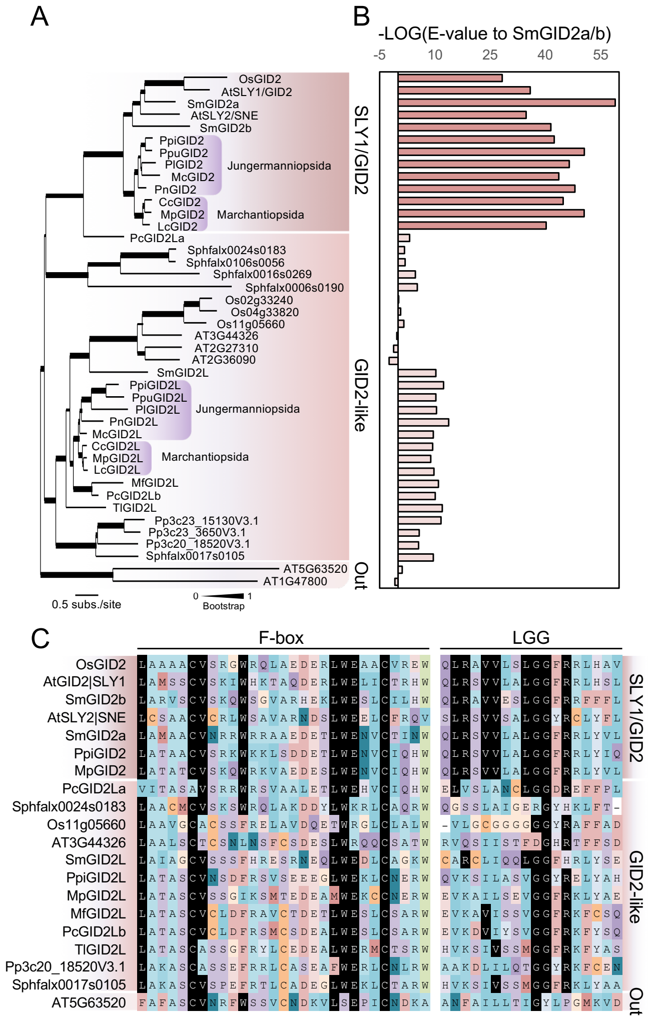


**Supplemental Figure 2. Presence of SLY1/GID2 orthologs in vascular plants**. (*A*) Phylogenetic analysis of SLY1/GID2 and GID2-like proteins in land plant representatives. Liverwort sequences from either Jungermanniopsida and Marchantiopsida classes are pointed. Support values associated with branches and displayed as bar thickness are maximum likelihood bootstrap values from 1000 replicates. (*B*) BLASTP value obtained for SmGID2a or SmGID2b as result using SLY1/GID2 and GID2-like proteins as queries. Value shown as the –LOG of the E-value retrieved in the BLASTP search. (*C*) Alignment showing highly conserved regions within the F-box domain and the LGG domain of selected SLY1/GID2 and GID2L proteins from A and B. Black colored boxes represent conserved sites in all SLY1/GID2 proteins.

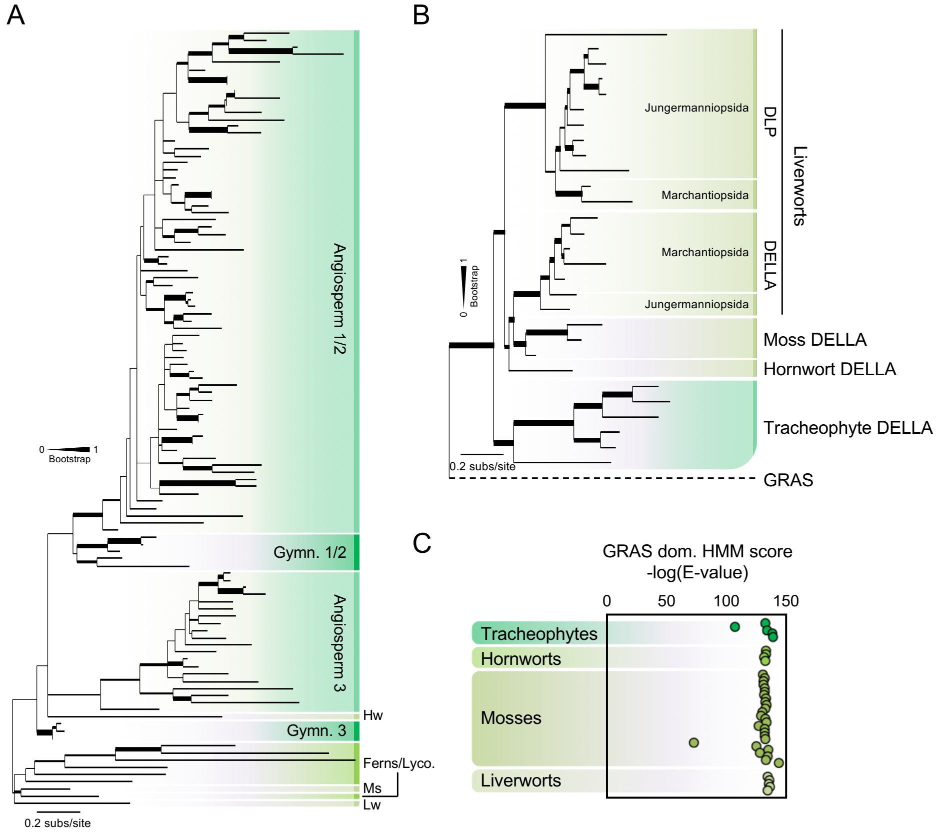


**Supplemental Figure 3. Non-vascular land plants have conserved DELLA proteins**. (*A*) Phylogenetic analysis of DELLA proteins using DELLA domain. (*B*) Phylogenetic analysis of liverwort DELLA and DELLA-like proteins using GRAS domains. Support values associated with branches and displayed as bar thickness are maximum likelihood bootstrap values from 1000 replicates. (*C*) Automated analysis of GRAS domain presence in non-vascular land plants using Pfam website. Scores are represented as the -Log of the E-value retrieved from the search and represented following the phylogenetic position obtained in Fig. 3B.

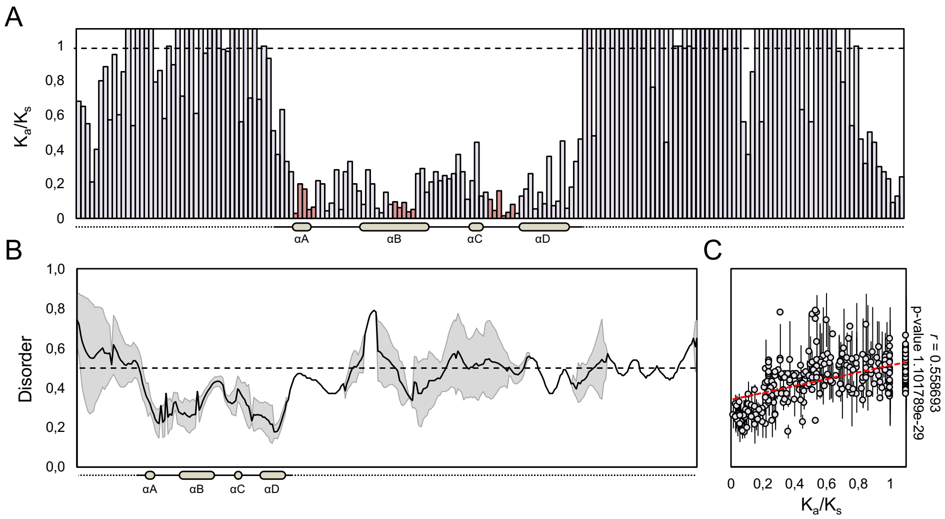


**Supplemental Figure 4. DELLA domain is structurally conserved**. (*A*) Selection analysis of the DELLA N-terminal región, based on codon-aligned nucleotide sequences. Non-conserved stretches generating gaps were removed. Red bars indicate residues of DELLA, LEQLE and TVHYNP motifs in the GA-conserved degron. Dashed line indicates the theoretical threshold for positive selection (above line) or purifying selection (below). (*B*) Analysis of disorder in several DELLA domains. Disorder values are plotted in a previously made alignment. Grey forms represent standard deviation from the average (black line). Dashed line indicates threshold value for ordered (<0.5) or disordered (>0.5) residues. (*C*) Relation between disorder and selection values per residue excluding gaps. p-value indicates Pearson's product moment correlation (*r*) statistical analysis. In general, less ordered stretches were subject to stronger positive selection.

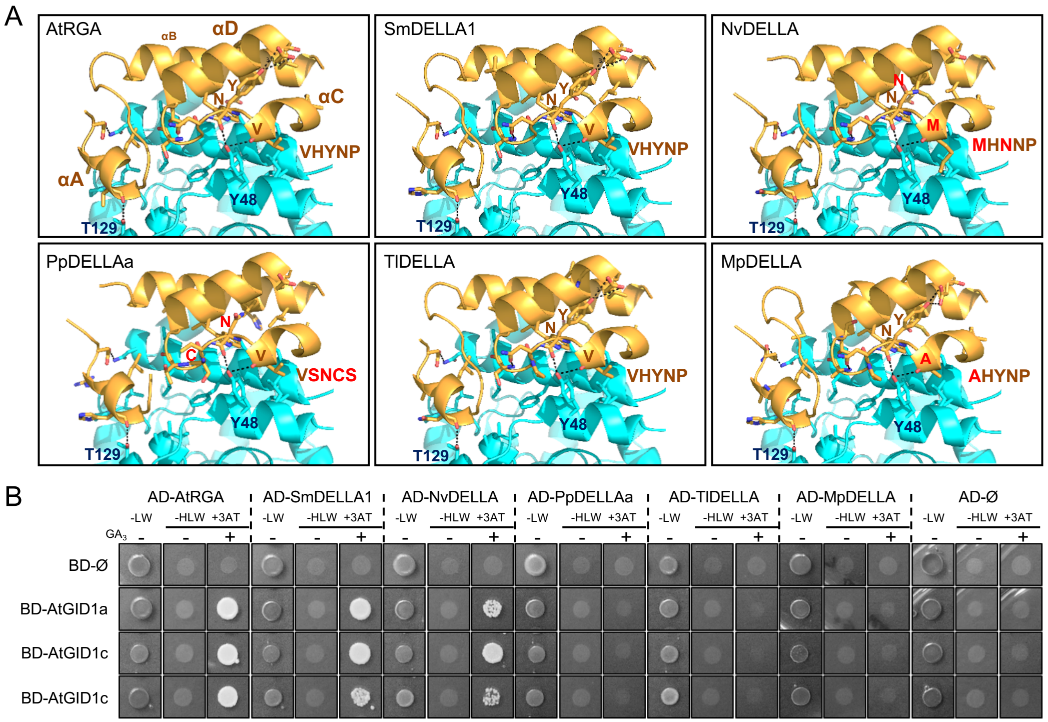


**Supplemental Figure 5. Some non-vascular DELLA proteins can interact with Arabidopsis GID1 receptors in a GA dependent manner**. (*A*) Predicted structural model for DELLA-AtGID1a interaction using AtRGA, SmDELLA1, NvDELLA, PpDELLAa, TlDELLA, and MpDELLA. DELLA structure is shown in yellow and AtGID1a structure in light blue. AtGID1a residues involved in DELLA interaction are written in dark blue. Residues different to that of AtRGA/GAI in main motifs are presented in red. Possible residue to residue interactions affected are pointed with a red circle. 90º rotation compared to Fig. 5A. (*B*) Yeast-two-hybrid assay between DELLA proteins and the three Arabidopsis GID1 receptors with or without 100 μM $GA_3$ (L, Leucine; W, Tryptophan; H, Histidine; 3-AT, 5 mM 3-aminotriazol). BD, GAL4 binding domain; AD, GAL4 activation domain.

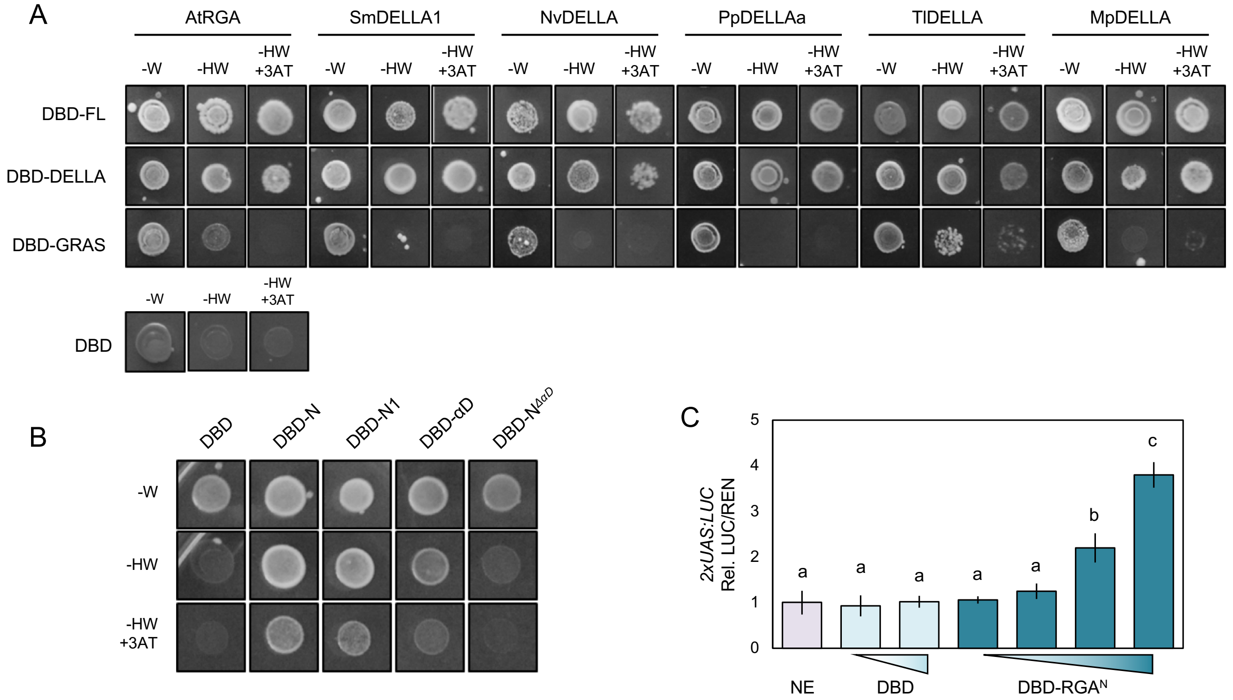


**Supplemental Figure 6. The conserved DELLA domain acts as a transcriptional activator domain**. (*A*) Yeast transactivation assay using DELLA protein full-length coding regions (DBD-FL), or truncated versions using either the GRAS domain (DBD-GRAS) or the DELLA domain (DBD-DELLA). (*B*) Yeast transactivation assay using different truncated versions of PpDELLAa DELLA domain as indicated in Figure 6C. (W, Tryptophan; H, Histidine; 3-AT, 5 mM 3-aminotriazol). BD, GAL4 binding domain; AD, GAL4 activation domain. (*C*) Dual luciferase transactivation assays in *Nicotiana benthamiana* leaves using the *LUC* gene under the control of the *Gal* operon UAS promoter as the reporter, and different effector vectors fused to the GAL4 DNA binding domain. NE, no effector; DBD, GAL4 binding domain; DBD-RGA$^{N}$, RGA DELLA domain fused to GAL4 DNA binding domain. Constitutively expressed *Renilla* luciferase (*REN*) for normalization. Data shown are normalized to NE value and represent the average of three biological replicates. Error bars represent standard deviation. Letters indicate significant differences between groups (P < 0.01, one-way ANOVA, Tukey's HSD post hoc test).

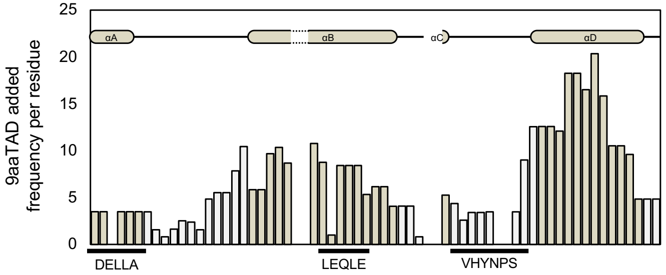


**Supplemental Figure 7. DELLA domain α helix D harbours a transactivation domain needed for recruitment of PolII co-activators**. Transactivation prediction plotted as the cumulative probabilities of 9aaTAD presence per residue found for AtRGA, SmDELLA1, NvDELLA, PpDELLAa, TlDELLA, and MpDELLA.
